# *Csf1rb* mutation uncouples two waves of microglia development in zebrafish

**DOI:** 10.1101/2020.11.04.368183

**Authors:** Giuliano Ferrero, Magali Miserocchi, Elodie Di Ruggiero, Valérie Wittamer

## Abstract

In vertebrates, the ontogeny of microglia, the resident macrophages of the central nervous system, initiates early during development from primitive macrophages. While murine embryonic microglia then persist through life, in zebrafish these cells are transient, as they are fully replaced by an adult population originating from larval hematopoietic stem cell (HSC)-derived progenitors. *Colony-stimulating factor receptor 1 (csf1r)* is a fundamental regulator of microglia ontogeny in vertebrates, including zebrafish which possess two paralogous genes: *csf1ra* and *csf1rb.* While previous work showed invalidation of both genes completely abrogates microglia development, the specific contribution of each paralog remains largely unknown. Here, using a fate-mapping strategy to discriminate between the two microglial waves, we uncover non-overlapping roles for *csf1ra* and *csf1rb* in hematopoiesis, and identified *csf1rb* as an essential regulator of adult microglia development. Notably, we demonstrate that *csf1rb* positively regulates HSC-derived myelopoiesis, resulting in macrophage deficiency, including microglia, in adult mutant animals. Overall, this study contributes to new insights into evolutionary aspects of Csf1r signaling and provides an unprecedented framework for the functional dissection of embryonic versus adult microglia *in vivo.*

## INTRODUCTION

Microglia are tissue-resident macrophages that play key immune and housekeeping roles in the central nervous system (CNS) (Prinz et al., 2019; Sierra et al., 2019). During development, microglia supports neurogenesis by releasing trophic factors (Tong and Vidyadaran, 2016), efficiently engulfing apoptotic neurons (Peri and Nusslein-Volhard, 2008) and pruning supernumerary synapses (Paolicelli et al., 2011). In the adult brain, microglia actively protrude branches to monitor the CNS microenvironment and interact with other cell types in order to maintain homeostasis (Davalos et al., 2005; Nimmerjahn et al., 2005). The indispensable role of microglia to foster CNS homeostasis becomes evident in human genetic conditions causing microglia deficits or dysfunctions (Li and Barres, 2018), which can result in severe pathologies such as Nasu-Hakola disease (Paloneva et al., 2002) and adult onset leukoencephalopathy with spheroids (Rademakers et al., 2011). Moreover, microglia are regarded as key mediators of the severe and prolonged inflammatory response triggered by CNS damage, which represents a major therapeutic hurdle in neurodegenerative disorders (Colonna and Butovsky, 2017).

During embryogenesis, microglia arise from yolk sac-derived primitive macrophages, which seed the developing neuroepithelium before the onset of neurogenesis (Alliot et al., 1999; Boche et al., 2012; Cuadros et al., 1993; Herbomel et al., 1999). Lineage-tracing studies performed in the mouse model showed these early microglia are maintained throughout life (Ginhoux et al., 2010), although later hematopoietic waves might partially contribute to the adult microglia pool (De et al., 2018). Similar to the mouse, microglia ontogeny in zebrafish initiates from amoeboid-shaped primitive macrophages, which colonize the neural tissue starting at 35 hours post-fertilization (hpf) and then differentiate into branched microglia at around 60 hpf (Herbomel et al., 2001). Unlike their mammalian counterparts however, embryo-derived microglia do not maintain in zebrafish, and the adult microglial network is established through a second wave of progenitors that seed the brain parenchyma later during development, fully replacing the initial population by the end of the juvenile stage (Ferrero et al., 2018; Xu et al., 2015). Cell transplantation and fate mapping experiments identified embryonic hematopoietic stem cells (HSCs) arising from the hemogenic endothelium in the dorsal aorta (DA), as the source of adult microglia in this model (Ferrero et al., 2018). Collectively, these findings have opened new avenues of research regarding possible functional differences between the two zebrafish microglial waves, as well as between mouse and zebrafish adult microglia, owing to their distinct cellular origins. Although zebrafish genetic models deficient for each microglia population would facilitate such comparative studies, little is known regarding the genetic regulation of adult microglia ontogeny, and so far no viable mutant resulting in the specific loss of adult microglia has been reported.

The tyrosine kinase *colony-stimulating factor receptor 1 (csf1r),* also known as M-csfr, is a fundamental regulator of mononuclear phagocyte homeostasis in vertebrates (Stanley and Chitu, 2014). It is predominantly expressed in macrophages and their precursors (regardless of their developmental origin), and exhibits pleiotropic effects including cell proliferation, differentiation and survival. Accordingly, CSF1R deficiency in human and mouse leads to a dramatic reduction in tissue macrophage development, including microglia (Erblich et al., 2011; Oosterhof et al., 2018; Rojo et al., 2019). Once established, microglia also rely on CSF1R signaling for their maintenance in the brain parenchyma and can be efficiently depleted in the mouse brain through pharmacological blockade of CSF1R (Elmore et al., 2014; Squarzoni et al., 2014). In humans, deficiencies in CSF1R signaling have been associated to neurodegenerative disorders, further highlighting the central role of CSF1R in microglia homeostasis (Oosterhof et al., 2019; Rademakers et al., 2011). *In vivo,* two non-homologous cytokines serve as ligands for CSF1R: Csf-1 and Interleukin 34 (II-34) (Lin et al., 2008; Stanley, 1977). Both show distinct spatial and cellular distribution in the brain parenchyma (Cahoy et al., 2008; Zeisel, 2015) and elicit both overlapping and non-redundant biological responses in regional microglia (Easley-Neal et al., 2019; Greter et al., 2012; Kana et al., 2019; Wang et al., 2012).

As a result of a teleost-specific whole genome duplication, zebrafish possess two paralogs of the *csf1r* gene: *csf1ra* and *csf1rb* (Braasch et al., 2006). Fish deficient in both genes *(csf1r^DM^)* lack microglia from the embryonic to the adult stages (Oosterhof et al., 2018), thus mimicking the *Csf1r^-/-^* mouse phenotype (Dai, 2002; Ginhoux et al., 2010). In contrast, individual mutants exhibit a less severe microglial phenotype, characterized by a transient loss of microglia in *csf1ra^-/-^* zebrafish embryos and a moderate reduction of adult microglia in both *csf1ra^-/-^* and *csf1rb^-/-^* single mutants (Oosterhof et al., 2018). Based on these phenotypes, it was suggested that both paralogs exhibit redundant functions. This prompted us to revisit the precise contribution of each paralog to microglia ontogeny, in light of the newly established model of microglia ontogeny in zebrafish where the two distinct primitive and definitive microglia populations temporally overlap. We previously demonstrated that the kdrl:Cre model offers a powerful tool to discriminate primitive macrophage-derived embryonic microglia from HSC-derived adult microglia *in vivo* (Ferrero et al., 2018). Exploiting this approach, we uncovered non-overlapping functions for *csf1ra* and *csf1rb* and identified *csf1rb* as a unique regulator of adult microglia development. In addition, we also demonstrated a specific contribution for the *csf1rb* paralogue to HSC-derived myelopoiesis, consistent with the specific HSC origin of the adult microglial population.

## RESULTS

### *Csf1rb* localizes to definitive hematopoiesis and to embryonic microglia during embryogenesis

Previous studies have documented the restricted expression of *csf1ra* in neural crest-derived cells, early macrophages and microglia during embryonic development (Caetano-Lopes et al., 2020; Herbomel et al., 2001; Parichy DM, 2000) (Fig. 1A-E), as well as its role in primitive myelopoiesis (Herbomel et al., 2001). However, although *csf1rb* has been previously linked to microglia biology (Mazzolini et al., 2019; Oosterhof et al., 2018), its expression has not been assessed in the context of developmental hematopoiesis and little is known about its specific functions. Using whole *in situ* hybridization (WISH), we found that *csf1rb* exhibits an expression profile distinct from that of *csf1ra* during embryogenesis, with no expression in either neural crest or primitive macrophages. Rather, *csf1rb* transcripts are first detected at around 30 hpf in the otic vesicle, as well as in a small number of hematopoietic cells in the posterior blood island (PBI) (Fig. 1F). The latter is consistent with expression in erythro-myeloid progenitors (EMPs), which are transiently found in the developing embryo. At 36 hpf, expression of *csf1rb* increases in the otic vesicle, and appears in cells located along the dorsal aorta (DA), a site at the onset of hematopoietic stem cell (HSC) formation (Fig. 1G). Over subsequent stages, expression in the otic vesicle disappears but expands in the DA and at 48 hpf onwards, *csf1rb* expression is observed in the caudal hematopoietic tissue (CHT), region of definitive hematopoiesis (Fig. 1H). At 72 hpf, the developing thymus also contains *csf1rb*-expressing cells, reminiscent of HSC-derived lymphoid progenitor immigrants (Fig. 1I, J). Notably, although *csf1ra* and *csf1rb* show distinct spatial expression patterns during development, both transcripts overlap in microglia in the brain and retina starting at 72 hpf (Fig. 1E, J).

**Figure 1.**
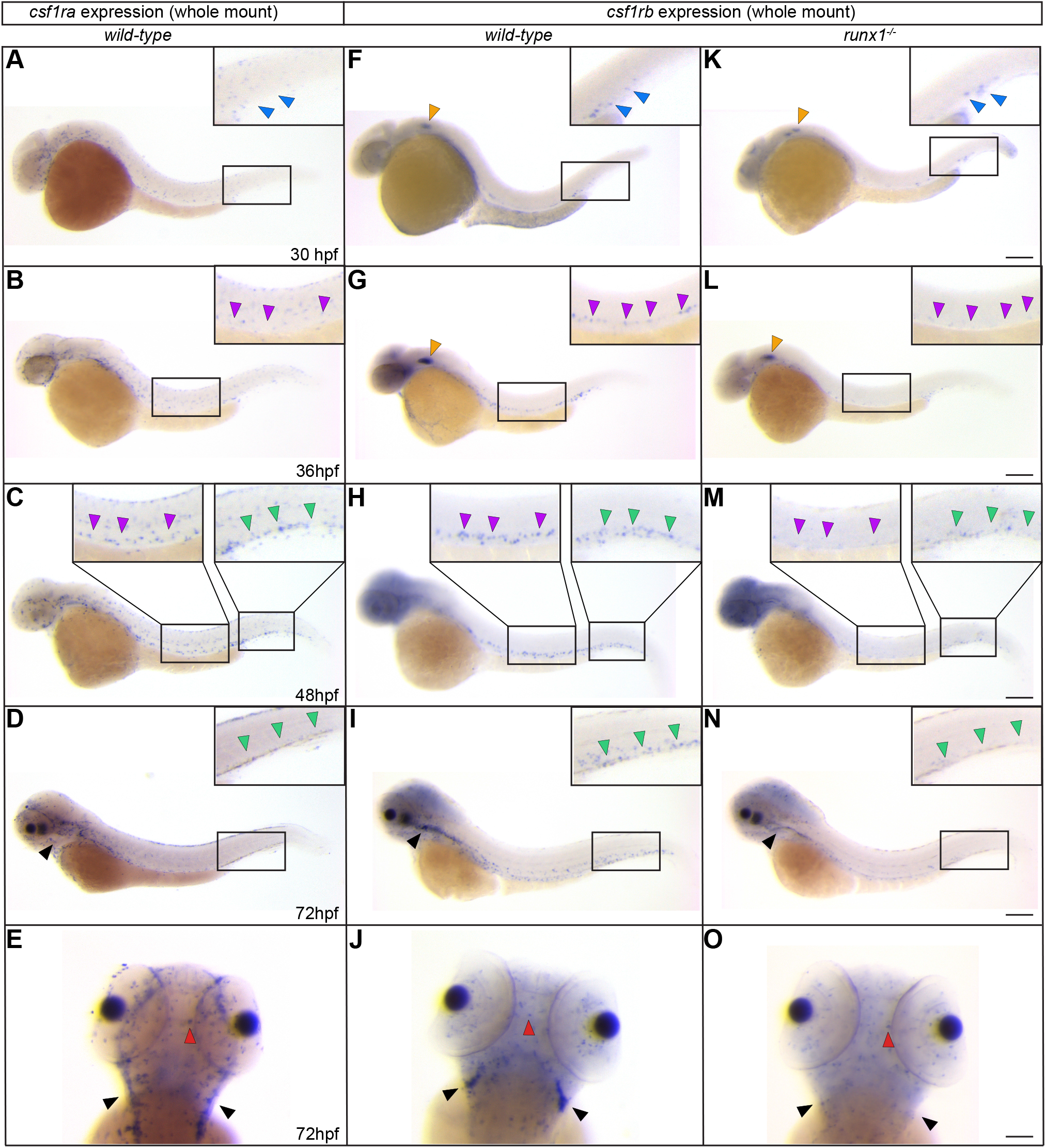
*Csf1ra* and *csf1rb* paralogs have nonoverlapping distribution during early development, except for microglia. Whole-mount in-situ hybridization (WISH) expression profiles of *csf1ra* (A-E) and *csf1rb* (F-J) in *wild-type* and *csf1rb* in *runx^-/-^* (K-O) embryos, at the indicated stages. All lateral views, except for E, J and O, shown in dorsal view. Orange and blue arrowheads indicate expression in the otic vesicle and posterior blood island (PBI) region, respectively. Purple and green arrowheads indicate expression in the dorsal aorta (DA) and caudal hematopoietic tissue (CHT), respectively. Black arrowheads show bilateral thymi and red arrowheads are microglial cells. Scale bars: 200 μm (30 and 36 hpf); 180 μm (48 hpf); 150 μm (72 hpf).

Because expression of *csf1rb* was mainly found in sites of definitive hematopoiesis, we performed WISH in *runx1* mutant embryos, which lack HSCs. While expression in the otic vesicle and in the PBI at 30 hpf is normal, we observed a strong reduction of *csf1rb* transcripts in the DA, CHT and thymus of homozygous embryos, thus identifying *csf1rb*-expressing cells in these anatomical locations as HSC-dependent (Fig. 1K-O). As expected, microglial expression of *csf1rb* was not affected in *runx1^nul1^* embryos, consistent with their ontogenic relationship with primitive macrophages, which are *runx1*-independent (Ferrero et al., 2018) (Fig. 10). Collectively, these results indicate that *csf1ra* and *csf1rb* paralogs have nonoverlapping distribution during early development, except for microglia.

### *Csf1rb* expression is restricted to hematopoietic progenitor cells and microglia among adult mononuclear phagocytes

We next assessed transcript expression of *csf1r* paralogs in mononuclear phagocytes isolated from adult tissues. As a source for these studies, we used *Tg(mhc2dab:GFP; cd45:DsRed)* double transgenic fish, as we previously demonstrated that the *mhc2dab:GFP; cd45:DsRed* transgene combination enabled the isolation by FACS of pure populations of tissue macrophages (Wittamer et al., 2011), including resident microglia (Ferrero et al., 2018). As shown in Fig. 2A, *csf1ra* was highly expressed in adult microglia, as well as in mononuclear phagocytes isolated from whole kidney marrow (WKM), spleen, liver and skin. In contrast to the ubiquitous expression pattern of *csf1ra,* we found high levels of *csf1rb* transcripts in microglial cells, very little expression in WKM macrophages and no expression in skin and spleen macrophages. Analysis of a publicly available WKM single cell dataset (Lareau et al., 2017) confirmed the lower expression of *csf1rb* versus *csf1ra* in macrophages, but also revealed major differences, with *csf1rb* being found specifically enriched within hematopoietic progenitor cells (Fig. 2B-E). Together with our WISH analyses, these results suggest that *csf1rb* expression labels blood progenitors through life and identified microglia as a unique population of mononuclear phagocytes to display *csf1rb* expression outside the WKM.

**Figure 2.**
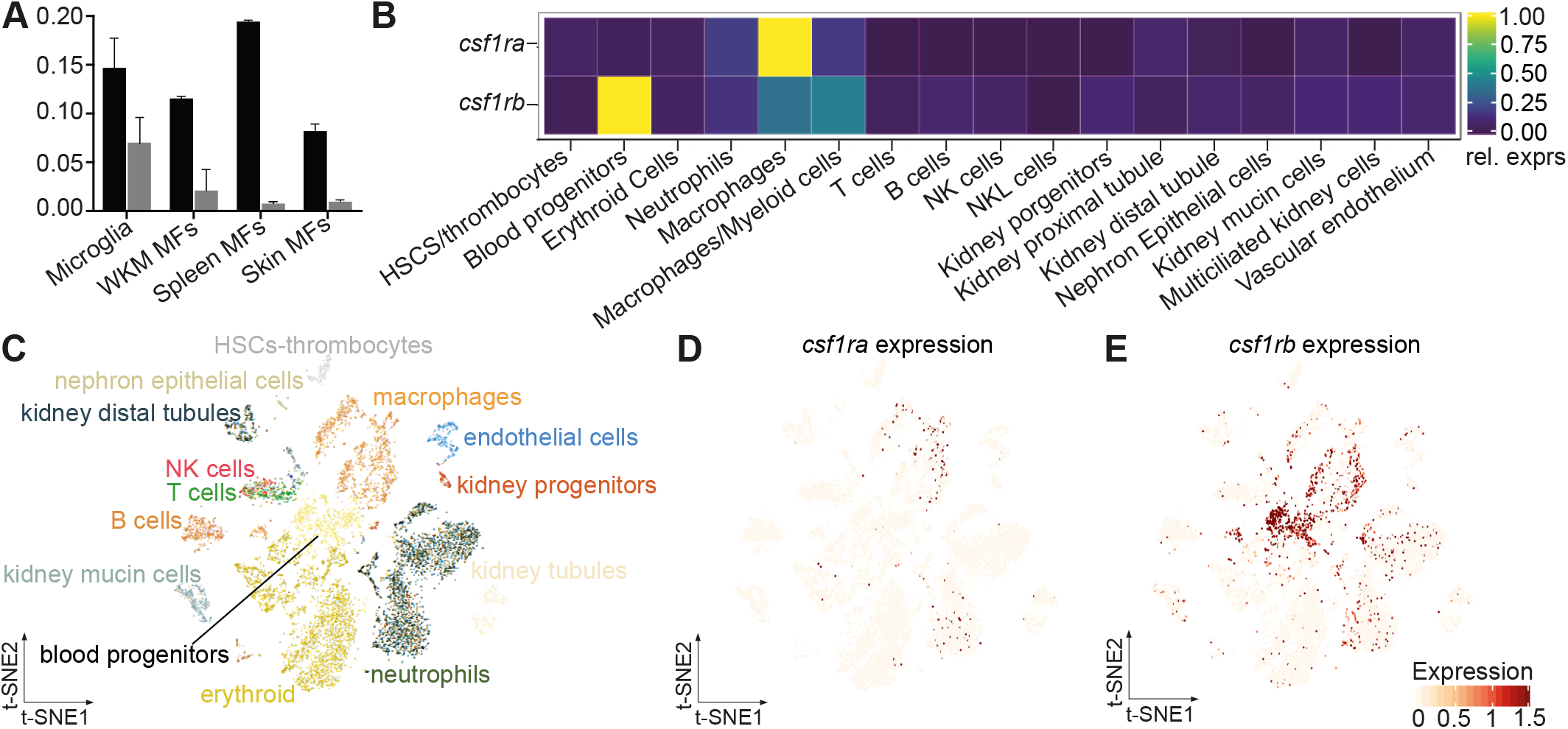
Characterization of *csf1r* paralog expression in adult hematopoietic cells. (A) QPCR expression for *csf1ra* and *csf1rb* in *cd45:DsRed^+^; mhc2dab:GFP^+^* mononuclear phagocytes sorted from adult zebrafish organs. Values on the y-axis indicate transcript expression normalized to *eflα* expression level. Error bars represent SEM (n=3). WKM: whole kidney marrow, MF: macrophages. (B-E) Expression profiles of *csf1r* paralogs in adult WKM hematopoietic and non-hematopoietic populations by single-cell RNAseq analysis, extracted from the public database from Laureau et al, 2017. Heatmap (B), 2D projection of the t-SNE analysis showing the distinct clusters identified in the adult WKM (C) and profiles of *csf1ra* (D) and *csf1rb* (E) across the clusters of the tSNE plot. Intensity of the color is proportional to the expression level.

### Different roles of *csf1ra* and *csf1rb* during embryonic microglia development

To study *csf1r* function *in vivo,* we used two zebrafish mutant lines with no functional *csf1ra* or *csf1rb* paralog. The zebrafish *panther* line carries a point mutation in *csf1ra,* replacing a valine by a methionine in position 614 (Parichy DM, 2000). This change induces an impaired functioning of the kinase activity of the receptor, resulting in the disruption of internal cell signaling. This model has been previously used to demonstrate the contribution of *csf1ra* to microglia development (Herbomel et al., 2001; Oosterhof et al., 2018) and constitutes therefore a valuable tool for our investigations. The zebrafish line *sa1503* harbors a splice site mutation in the *csf1rb* gene, leading to the inclusion of 86 nucleotides from intron 11 and a premature stop codon (Fig. S1A). This nonsense mutation results into the synthesis of a truncated protein that lacks the receptor kinase domain and is expected to be non-functional. The presence of the non-spliced transcript was confirmed through RT-PCR and sequencing analyses, thus validating *csf1rb* loss-of-function in the mutant (Fig. S1B, C). Homozygous *csf1rb*^sa1503^ fish exhibit normal external morphology and behavior and, like the *csf1ra^-/-^* mutant, survive to adulthood.

Previous studies indicated that *csf1ra* is not required for early myelopoiesis, as primitive macrophages develop normally in *csf1ra^-/-^* embryos (Herbomel et al., 2001). Extending these analyses, we found no effect on the number of *mfap4^+^* primitive macrophages in *csf1rb* mutants, as determined using whole-mount *in situ* hybridization (Fig. 3A,B). To study whether both paralogs were simultaneously required for primitive myelopoiesis, we intercrossed the two single mutant lines and derived *csf1ra/b* double mutant embryos (hereafter referred to as *csf1r^DM^).* As shown in Fig. 3A,B, the complete loss of *csf1r* had no consequence on primitive macrophage ontogeny, as the number of *mfap4^+^* cells in double mutants were similar to that of *wildtype* and single homozygous mutant embryos. This is consistent with our recent findings using the macrophage *mpeg1:EGFP* reporter line (Kuil et al., 2020). We next investigated the requirement of the different *csf1r* paralogs for the establishment of embryonic microglia, which differentiate in the brain parenchyma from primitive macrophages starting at 60 hpf (Ferrero et al., 2018; Herbomel et al., 2001). As readout for microglia differentiation, we analyzed by WISH the expression of *apoeb,* a microglia signature gene. Quantification of *apoeb^+^* cells present in the optic tectum at 72 hpf showed the number of microglia was dramatically decreased in *csf1ra*-deficient embryos (0.8 ± 0.4 cells) when compared to *wild-type* (20.8 ± 1.4 cells), unaffected in *csf1rb*-depleted embryos (19.1 ± 1.3 cells) and similarly strongly reduced in *csf1r^DM^* embryos (0.8 ± 0.5 cells) (Fig. 3C,D). These results indicate that independently, *csf1ra* and not *csf1rb,* is important for establishing the first wave of microglia during zebrafish embryogenesis.

**Figure 3.**
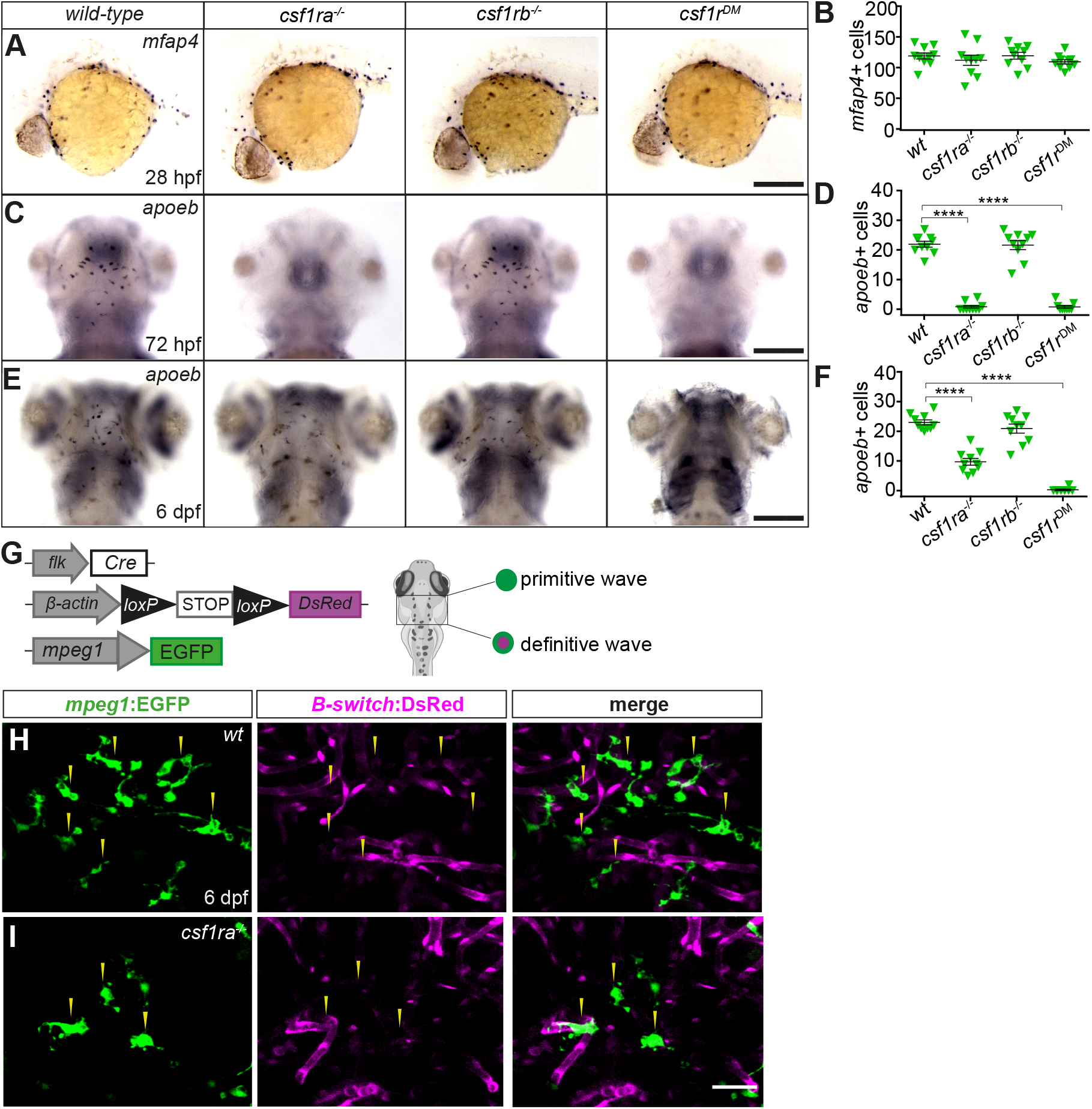
*csf1ra* and *csf1rb* are differently required for embryonic microglia development. (A, C, E) WISH of the indicated genes in *wild-type, csf1ra^-/-^, csf1rb^-/-^* and *csf1r^DM^* siblings, at the stages indicated. All dorsal views except for A, shown in lateral view. Scale bars: 150 μm (A, C); 100 μm (E). (B, D, F) Quantification of *mfap4^+^* primitive macrophages (B), *apoeb^+^* microglia at 3 dpf (D) and at 6 dpf (F) in the indicated genotypes. Each symbol represents a single embryo/ larvae and error bars represent mean ± SEM. Differences between groups were analyzed by Students t-test [***p<0.001; ****p<0.0001]. (G) Scheme of the transgenic lines used to discriminate the primitive and definitive microglia waves in 6 dpf larvae. (H, I) Imaging by confocal microscopy of the optic tectum in 6 dpf wild-type (H) and *csf1ra^-/-^* (I) sibling larvae carrying the *kdrl:Cre; ßactin:Switch-DsRed; mpeg1:EGFP* triple transgene. GFP (left panels), DsRed (middle panels) and merge of both fluorescence channels (right panels) are shown. Images were taken with an inverted Zeiss LM780 confocal microscope using a 25x water-immersion objective. Scale bar: 50μm.

Because it was previously reported that *csf1ra^-/-^* embryos exhibit a partial recovery of microglia cells at 6 dpf (Herbomel et al., 2001), we next examined the status of microglia in the mutants later during development using WISH. While the numbers of *apoeb+* cells were stable from 3 to 6 dpf in *wild-type* and *csf1rb^-/-^* embryos (approximately 20 cells per optic tectum), we observed a gradual increase in the number of microglia (from 0.7 ± 0.4 cells at 3 dpf to 9.7 ± 1.1 cells at 6 dpf) in embryos carrying the *csf1ra^-/-^* mutation (Fig. 3E,F). At 6 dpf microglia cell numbers in *csf1ra^-/-^* embryos accounted for approximately 50% of total microglia cells found in sibling controls. Interestingly, at the same developmental stage, repopulation of the brain parenchyma by microglia was not observed in double mutant embryos, which remained devoid of *apoeb*-expressing cells. This observation indicated that recovery of microglia in *csf1ra*-deficient embryos is mediated by *csf1rb,* which suggested there may be a compensatory role for *csf1rb* in microglia development in the absence of *csf1ra.* However, when we FACS-sorted *mpeg1:EGFP+* cells from the heads of 6 dpf *wild-type* and *csf1ra^-/-^* embryos, we found no significant difference in expression of *csf1rb* transcripts between both genotypes (Fig. S2). Taken together, these data suggest that the partial recovery of embryonic microglia in *csf1ra^-/-^* embryos is *csf1rb-* dependent but does not require a compensatory increase in *csf1rb* mRNA.

We investigated the source of the repopulating microglial cells in *csf1ra*-deficient embryos. Indeed, microglia recovery in these embryos could result either from a delay of differentiation of primitive macrophages or from the early and atypical contribution of HSCs, the precursors of adult microglia. We discriminated between these two possibilities by crossing the *csf1ra* mutant line to *Tg(kdrl:Cre; bactin2:loxP-Stop-loxP-DsRed^express^* (also known as ßactin:Switch-DsRed); *mpeg1:EGFP)* triple transgenics (Fig. 3G). As we previously showed, primitive macrophage-derived embryonic microglia are GFP^+^, DsRed^-^ in this setup (Fig. 3H), while mononuclear phagocytes originating from EMPs or HSCs are GFP^+^, DsRed^+^, owing to the hemogenic nature of their precursors (Ferrero et al., 2018). Confocal microscopy analysis of live embryos revealed that GFP^+^ microglia present at 6 dpf in *csf1ra-*deficient embryos did not express the DsRed transgene, thus demonstrating their lineage relationship with primitive macrophages (Fig. 3I). These findings indicate that recovered microglial cells in the *csf1ra* mutant share the same cellular origin as their *wild-type* counterparts and point to a delay of primitive macrophage differentiation as the cause of the observed phenotype.

### *csf1rb* is a regulator of definitive microglia

Like the single homozygous mutants, *csf1r^DM^* are viable and fertile, allowing for investigations into the role of Csf1r signaling in the establishment of definitive microglia. As a way to discriminate between embryonic and adult microglia in our analyses, we relied again on mutant fish carrying the *kdrl*:Cre; ßactin:Switch-DsRed; *mpeg1:EGFP* triple transgene (Fig. 4A) and performed confocal analyses of brain sections immuno-stained for GFP and DsRed. In line with previous findings, the density of GFP^+^ microglia cells in the brain parenchyma was decreased ~60% in single *csf1ra^-/-^* and *csf1rb^-/-^* mutant fish as compared to their *wild-type* siblings (Fig. C,D,N) and showed a dramatic reduction (90%) in adult animals lacking both paralogs (Fig. 4E,N). However, analyzes for DsRed transgene expression to assess their primitive or definitive identity revealed striking microglial phenotypes (Fig. 4F-O). In *wild-type* fish, all *mpeg1^+^* microglia were DsRed^+^, as expected from their known HSC origin (Fig. 4 F,J,O). Similarly, GFP^+^ microglial cells from *csf1ra^-/-^* fish also co-expressed DsRed, indicating that adult microglia ontogeny still occurs in the absence of *csf1ra* (Fig. 4 G,K,O). In contrast, the majority of the remaining GFP^+^ cells in *csf1rb^-/-^* animals were found to be DsRed^-^, thus excluding them as microglia derived from the adult wave (Fig.4 H,L,O). Based on the lack of DsRed expression, these mpeg1:EGFP^+^ cells likely represent residual primitive microglia. This is further supported by observations that in *csf1r^DM^* animals, which lack primitive microglia, the very few cells present in the brain parenchyma all expressed DsRed (Figure 4 I,M,O). Collectively, these data indicate that *csf1rb,* and not *csf1ra,* is essential for establishing the definitive wave of microglia in zebrafish.

**Figure 4.**
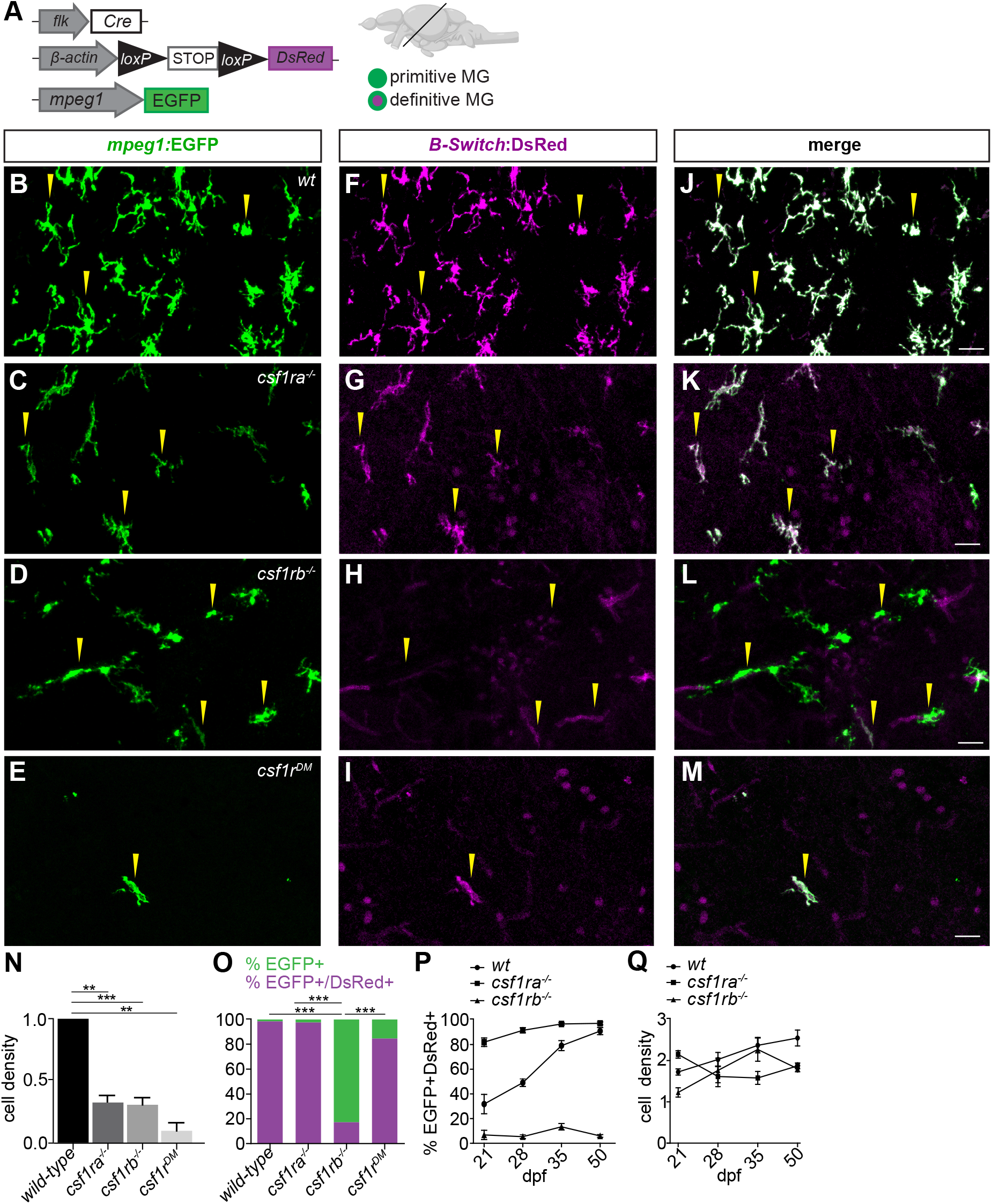
*csf1rb* is required for HSCs-derived microglia development. (A) Scheme of the transgenic lines used to discriminate primitive from definitive microglia in adult zebrafish brains. (B-M) Immunofluorescence on transversal brain sections from *Tg(kdrl:Cre; ßactin:Switch-DsRed; mpegl:EGFP)* triple transgenic adult *wild-type* (B, F, J), *csf1ra^-/-^* (C, G, K), *csf1rb^-/-^* (D, H, L) and *csf1r^DM^* (E, I, M) fish. GFP (left panels), DsRed (middle panels) and merge of both fluorescence channels (right panels) are shown. (N,O) Quantification of microglia density (GFP ^+^ cells/100 μm2) (N) and percentage of GFP^+^ DsRed^-^ (green) embryonic versus GFP^+^ DsRed+ (purple) adult microglia (O) in each genotype. Bars represent the mean ± SEM (n=4). For each individual, cells were counted on ten, 3Oμm-thick brain sections from rostral to caudal. Differences between groups were analyzed by One-way ANOVA test [***p<0.0005.]. Images were taken with an inverted Zeiss LM780 confocal using a 20x objective. Scale bars= 50μm. (P,Q) Percentage of GFP^+^ DsRed^+^ adult microglia (P) and microglia density (cells/1 mm3) (Q) at 21,28, 35 and 50 dpf for each genotype. Cells were counted on tissue-cleared whole brains as described in *Ferrero et al*, 2018. Cell counts were limited to the optic tectum and hindbrain areas. Between 4 (21 dpf, 28 dpf) and 6 (50 dpf) images per brain were acquired, with an average z-stack of 400 μm., using a Zeiss LM780 confocal with a 25x water-immersion objective.

To characterize the developmental dynamics leading to the observed phenotype, we examined the brains of *Tg(kdrl:Cre;* ßactin:Switch-DsRed; *mpeg1:EGFP) wild-type* and *csf1ra* or *csf1rb* mutants larvae at 21, 28, 35 and 50 dpf. As we previously reported, this time window encompasses the progressive replacement of GFP^+^ DsRed^-^ primitive microglia by definitive GFP^+^ DsRed^+^ microglia in the brain parenchyma (Ferrero et al., 2018). These kinetic analyses revealed distinct phenotypes among the mutants. Consistent with our previous observations, in *wildtype* fish the percentage of adult DsRed^+^ microglia steadily increased over time (Fig. 4P). In contrast, the brain of *csf1rb* mutants remained largely devoid of DsRed^+^ cells at all time points, suggesting that microglial progenitors fail to colonize the CNS in the absence of *csf1rb* (Fig. 4P). Surprisingly, in *csf1ra^nul1^* animals we observed a shift in the emergence of adult microglia. At 21 dpf, when GFP^+^ DsRed^-^ primitive microglia are still predominant in *wild-type* brains, the majority of microglia in *csf1ra^-/-^* fish already express DsRed (Fig. 4P). Based on these observations, we hypothesized that primitive microglia detected in the *csf1ra^-/-^* brain at 6 dpf fail to maintain through the juvenile stage. Interestingly, considering the overall density of mpeg1^+^ microglia across time, irrespective of the origin, we observed that in *wild-type* individuals the density of microglia increased from 1,7 ± 0.07 cells/mm^3^ to 2,5 ± 0.2 cells/mm^3^ between 21 and 50 dpf, mirroring the progressive expansion of the DsRed^+^ cells. In *csf1rb* mutants, microglia similarly expanded from 1,2 ± 0.1 to 2,2 ± 0.3 cells/mm^3^ between 21 and 35 dpf (Fig. 4Q). Given that these cells are from embryonic origin, such findings suggest that a partial compensation takes place in the brain of *csf1rb^-/-^* fish in the absence of DsRed^+^ adult microglia. However, the potential of primitive microglia to compensate for the lack of the adult wave appears to be limited, since cell density dropped to 1,8 ± 0.07 cells/mm^3^ at 50 dpf (Fig. 4Q) and remained lower than in *wild-type* fish throughout adulthood (Fig. 4N). The curve of microglia density across time followed a different trend in *csf1ra^-/-^* fish, where DsRed^+^ cells successfully established in the brain by 21 dpf, but did not undergo the steady expansion that we observed in *wild-type.* This result suggests that, while dispensable for their ontogeny, *csf1ra* is likely required for maintaining adult microglia after they colonize the juvenile brain. Overall, we concluded that the *csf1ra* and *csf1rb* paralogs are respectively required for the maintenance and specification of embryonic and adult microglia in zebrafish and that individual loss of function of either paralogue results in reduced microglia densities in the adult.

### *csf1rb* is required for the development of HSCs-derived myeloid cells

We sought to dissect the mechanisms linking *csf1rb* to adult microglia development. Based on the expression profile of *csf1rb* in hematopoietic progenitors and the developmental relationship between adult microglia and HSCs, we hypothesized that *csf1rb* regulates definitive hematopoiesis in zebrafish. By WISH, we did not detect any significant alteration in the expression of *runx1* in the DA of *csf1rb-* deficient embryos, indicating normal specification of the hemogenic endothelium (Fig. 5A,B). In addition, at 3 and 6 dpf, *c-myb* expression in the CHT and the pronephros, which specifically labels HSCs and progenitors, was not changed in *csf1rb^-/-^* embryos (Fig. 5C-F). This demonstrates that neither the emergence of HSCs nor the maintenance of progenitors during embryonic development requires *csf1rb.*

**Figure 5.**
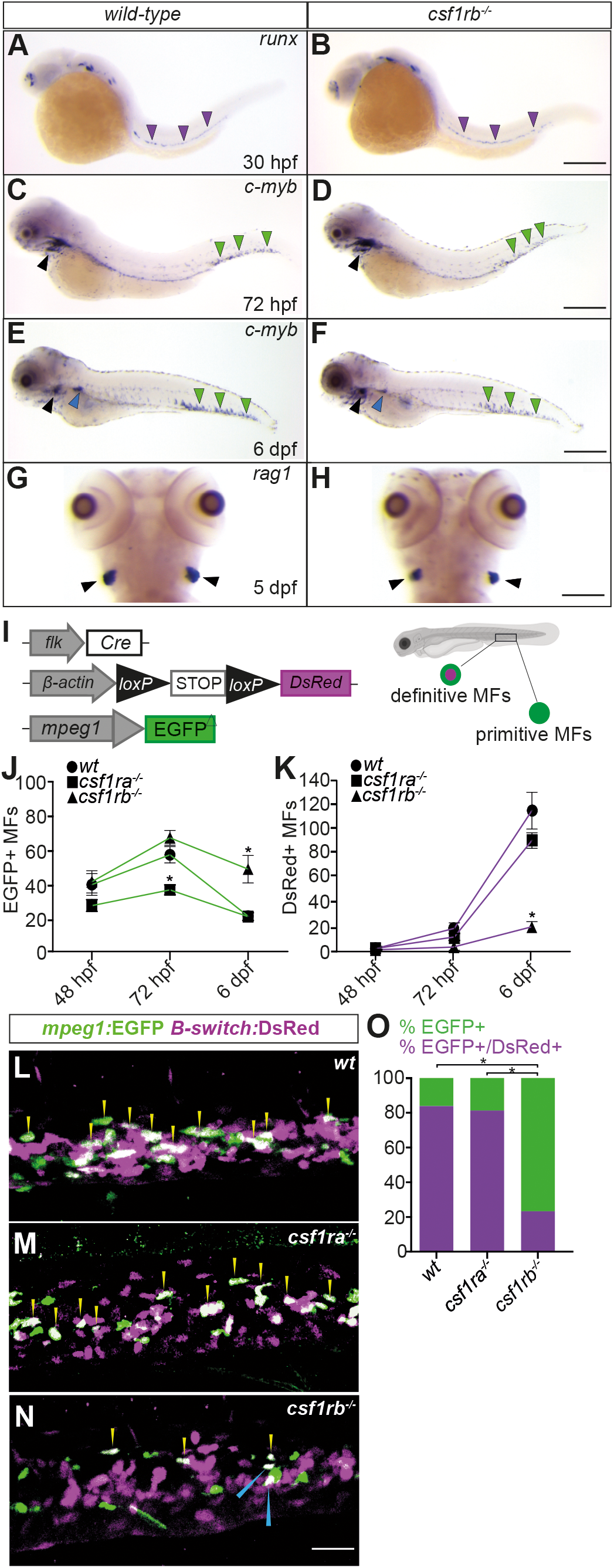
Loss of *csf1rb* function impairs the development of definitive macrophages. (A,H) WISH of the indicated genes in *wild-type* and *csf1rb^-/-^* siblings at the stages indicated. All lateral views except for G and H, shown in dorsal. The purple arrowheads indicate *runx1^+^* HSCs along the dorsal aorta. The green and blue arrowheads show *cmyb^+^* hematopoietic progenitors in the CHT and in the pronephros, respectively. Black arrowheads indicate *rag1^+^* or *cmyb^+^* lymphoid progenitors in the thymus. Scale bar: 200 μm (A, B), 150 μm (C, D, G, H), 100 μm (E, F). (I-O) Confocal imaging analysis of definitive myelopoiesis in *csf1r* mutant embryos and larvae (I) Scheme of the transgenic lines used to discriminate the primitive and definitive myelopoiesis waves in the CHT. (J,K) Quantification of GFP^+^ DsRed^-^ primitive (J) and GFP^+^ DsRed^+^ definitive macrophages (K) in *wild-type, csf1ra^-/-^* and *csf1rb^-/-^* carrying the *kdrl:Cre; ßactin:Switch-DsRed; mpeg1:EGFP* triple transgene, at the indicated developmental stages (n=4; symbols represent mean ± SEM). (L-N) Confocal imaging and quantification of the percentage (O) of GFP^+^ DsRed^-^ primitive macrophages (green) versus GFP^+^ DsRed^+^ definitive macrophages (purple) in the CHT of 6 dpf *wild-type* and *csf1r* mutant larvae. Bars in the graph represent the mean of 4 larvae for each group. Cells in the CHT were quantified in four contiguous 385 μm2 fields per CHT, with an average 100 μm z-stack, from caudal to rostral. Images were taken with an inverted Zeiss LM780 confocal microscope, using a 25x water-immersion objective. Difference between groups were analyzed by Kruskal-Wallis test [*P<0.05].

We evaluated a possible requirement for *csf1rb* during HSC differentiation. In the zebrafish embryo, T lymphopoiesis starts at around 50 hpf, with HSC-derived thymocyte precursors migrating to the developing thymus (Hess and Boehm, 2012; Murayama et al., 2006). We found that T cell development was not affected in the absence of *csf1rb,* as expression of the early T cell marker *rag1* was detected in mutant embryos at levels similar to that seen in *wild-type* (Fig. 5G, H). Next, we assessed the myeloid potential of *csf1rb*-deficient HSCs. As the different waves of myelopoiesis temporally overlap during embryonic development, we used triple transgenic *kdrl*:Cre; ßactin:Switch-DsRed; *mpeg1:EGFP* embryos to discriminate in the CHT between newly born definitive macrophages (GFP^+^ DsRed^+^) and primitive macrophages (GFP^+^ DsRed^-^) having colonized the site from the periphery (Fig. 5I). In *wild-type* embryos, we found that for the first 48 hours of development, all GFP^+^ macrophages present in the CHT (40 cells on average) are derived from primitive hematopoiesis, as indicated by their lack of DsRed expression (Fig. 5J, K). The first definitive macrophages, identified as DsRed^*+*^ GFP^+^ cells, were detected in the CHT at around 48 hpf (~3 cells on average) (Fig. 5J, K). This population then slowly increased over time, and at 6 dpf, the CHT contained on average 113 double positive cells per embryo. By that stage, definitive macrophages in the CHT outnumbered primitive macrophages, accounting for up to 80% of the total GFP^+^ population. Having delineate the kinetics of differentiation of definitive macrophages in the CHT, we next performed similar quantification of macrophage numbers in single mutants. In *csf1ra*-deficient embryos, we found that both the kinetics of appearance and total number of DsRed^+^ GFP^+^ double positive cells were similar to *wild-types*, thus indicating that the developmental program of HSC-derived macrophages was not affected (Fig. 5K-O). In contrast, *csf1rb^-/-^* embryos exhibited significantly decreased numbers of definitive macrophages at each time points (4 versus 19 at 72 hpf and 19 versus 113 at 6 dpf) (Fig. 5K-O). These findings demonstrate that Csf1rb activity specifically supports the embryonic development of HSC-derived macrophages. Interestingly, analysis of CHT GFP^+^ primitive macrophages during the 48 hpf to 6 dpf time-window also revealed major phenotypic differences among the mutants. Consistent with previous observations (Herbomel et al., 2001) and our own results suggesting a role for *csf1ra* in controlling early macrophage invasion in embryonic tissues, we found that the CHT of *csf1ra* mutants became colonized by 30% less primitive macrophages (Fig. 5J). By comparison, the numbers of primitive macrophages present in the CHT of *csf1rb* mutant embryos were similar to that of *wild-types.* These findings suggest that in primitive macrophages the *csf1rb* paralogue is dispensable for cell migration. Collectively, these results demonstrate that *csf1rb,* and not *csf1ra,* is required for definitive myelopoiesis in the zebrafish embryo.

To evaluate whether Csf1rb functions are required for life, we examined WKM cell suspensions from *mpeg1:EGFP* transgenics by flow cytometry. These analyses revealed the relative percentage of mpeg1:EGFP^high^ cells, which identify macrophages in adult fish (Ferrero et al., 2020), was reduced from 1 ± 0.2% for *wild-type* to 0.4 ± 0.2% for *csf1rb^-/-^* animals, while *csf1ra^-/-^* fish exhibited an intermediate value (0.7 ± 0.1%) (Fig. 6A). Intriguingly, the mpeg1:EGFP^low^ population, which labels most IgM^+^ B lymphocytes (Ferrero et al., 2020) was also barely detectable in *csf1rb^-/-^* animals (Fig. 6B) (Mean ± SEM; *wild-type:* 13 ± 3.6%; *csf1ra^-/-^*: 10.9 ± 0.3%; *csf1rb^-/-^*: 1.6 ± 0.5%). *Csf1rb* mutants also displayed a light-scatter profile distinct from that of *wildtype* and *csf1ra^-/-^* fish, with a significant loss of the forward scatter (FSC)^hi^ side scatter (SSC)^hi^ myeloid gate (Mean ± SEM, n=3; *wild-type:* 27.5 ± 1.7%; *csf1ra^-/-^:* 26 ± 4.2%; *csf1rb^-/-^:* 16.3 ± 1.7%), as well as an increase of the FSC^hi^ SSC^l0^ progenitor fraction (Mean ± SEM; *wild-type:* 22.2 ± 1.8%; *csf1ra^-/-^*: 28.6 ± 5.8%; *csf1rb^-/-^*: 45.5 ± 1.6%) (Fig. 6B). As the myeloid fraction mostly contains mature neutrophils, these observations suggest a complete block at the myeloid progenitor stage. Finally, and in line with the change in mpeg1-expressing B lymphocytes, the relative percentage of FSC^l0^SSC^l0^ lymphoid cells was also impaired in *csf1rb^-/-^* animals (Mean ± SEM; *wild-type:* 36.2 ± 2.2%; *csf1ra^-/-^:* 34.3 ± 3.6%; *csf1rb^-/-^:* 21.7 ± 0.4%) (Fig. 6A). Collectively, these findings indicate the absence of *csf1rb* results in functional deficiencies in myelopoiesis in the adult, and a concomitant lack of mpeg1-expressing B lymphocytes.

**Figure 6.**
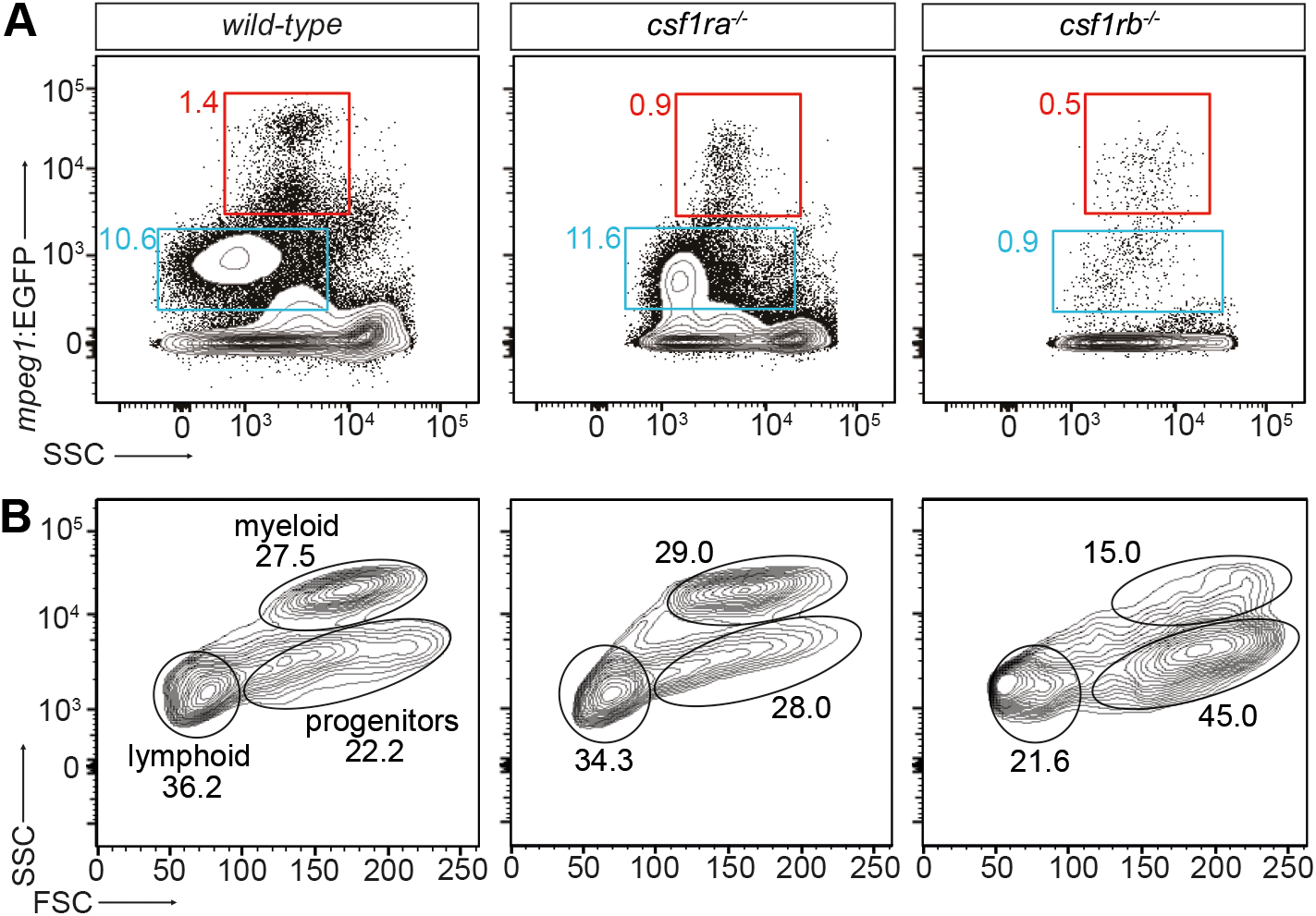
Adult zebrafish *csf1rb* mutant display hematopoietic deficiencies. (A, B) Flow cytometry analysis of WKM cell suspensions from *wild-type, csf1ra^-/-^* and *csf1rb^-/-^* adult fish carrying the *mpegl:EGFP* reporter. (A) The *mpeg1:EGFP^hi^* fractions identify mature macrophages (red frames), while the *mpeg1:EGFP^lo^* fractions contain mainly IgM-expressing B lymphocytes (blue frames). (B) Scatter profiles of WKM in typical *wild-type* (left panel), *csf1ra^-/-^* (middle panel) and *csf1rb^-/-^* (right panel) adult fish. Number in plots indicate percent of cells in circled myeloid, progenitor and lymphoid gates. Means ± SEM for 3 individuals are indicated in the text.

### DISCUSSION

Tissue macrophages constitute a highly heterogeneous compartment, based on their origin and the niche they inhabit (Bennett and Bennett, 2019; Guilliams et al., 2020). CSF1R signalling is a common pathway regulating the development of most macrophages in vertebrates, as demonstrated by their dramatic loss (including microglia) in *Csf1r*-deficient mice and *csf1r*-deficient zebrafish (Dai, 2002; Kuil et al., 2020; Oosterhof et al., 2018; Rojo et al., 2019). However, while *CSF1R* is represented only once in mammalian genomes, zebrafish possess two copies of the gene and their relative contribution to myelopoiesis has remained unknown. Focusing on the specific context of microglia ontogeny, in this study we have thus investigated the functions of each paralog during macrophage development. While our gene expression analyses demonstrate that *csf1rb,* like *csf1ra,* is expressed within the hematopoietic compartment, they also reveal a divergence in their expression profiles, with *csf1ra* expressed in all tissue macrophages (regardless of their primitive or definitive origin) and *csf1rb* restricted to microglia and definitive blood progenitors. Given that in the mouse *Csf1r* is expressed all throughout the path of macrophage differentiation (from hematopoietic progenitors to mature cells) (Hawley et al., 2018; Sasmono et al., 2003), such complementary patterns suggest that subfunctionalization, a process where the two gene copies partition the ancestral function (Force et al., 1999), may have contributed to the evolution of this family in zebrafish. Accordingly, *csf1ra* signalling is required for the establishment of primitive macrophage-derived embryonic microglia (Herbomel et al., 2001; Oosterhof et al., 2018), while *csf1rb* controls the ontogeny of definitive macrophages, including adult microglia. Our work thus demonstrates that *csf1ra* and *csf1rb* are jointly required to fulfil the roles of mammalian CSF1R in myelopoiesis.

Interestingly, despite these functional divergences, we also provide evidence that *csf1rb* is able to compensate, at least partially, for the absence of *csf1ra.* For example, whereas *csf1rb* loss has no effect on microglia development in embryos with a functional *csf1ra* paralog, *csf1rb* signalling is responsible for the partial recovery of microglia observed in *csf1ra^-/-^* larvae, as indicated by the absence of recovery in *csf1r^DM^* mutants. Since microglia repopulating *csf1ra^-1-^* larvae entirely derived from primitive macrophages and did not exhibit a compensatory overexpression of *csf1rb,* it thus appears that in this setting the basal expression level of *csf1rb* on embryonic microglia is sufficient for driving the recovery process. Our findings also provide new insights into the functions and complex interplay between csf1r paralogs and the three Csf1r ligands identified in zebrafish: Interleukin-34 (Il34), Csf1a and Csf1b. Similar to the mouse (Greter et al., 2012; Wang et al., 2012), Interleukin-34 acting through Csf1ra is thought to control the migration of primitive macrophages to the embryonic neuroepithelium in zebrafish (Kuil et al., 2019; Wu et al., 2018). Accordingly, *il34-* deficient embryos phenocopy both the microglial loss at 3 dpf and the partial recovery at 5 dpf observed in *csf1ra* mutants (Herbomel et al., 2001; Kuil et al., 2019). Given that *csf1r^DM^* larvae are completely devoid of microglia at 6 dpf, this suggests that *csf1rb*-mediated microglia replenishment in *csf1ra^-/-^* larvae is independent of Il34 signalling. Also, since the very few microglia present at 3 dpf in *csf1ra* or *il34*-deficient embryos retain their proliferative capacity and can be increased by *csf1a* overexpression (Kuil et al., 2019; Oosterhof et al., 2018), it is tempting to speculate that *csf1a* and/or *csf1b* signalling through *csf1rb* triggers the proliferation of embryonic microglia in *csf1ra* mutants. In line with this interpretation, we recently showed that primitive macrophages in *csf1r^DM^* embryos, which lose interaction with all Csf1r ligands, exhibit both migration and proliferation defects (Kuil et al., 2020). Further investigation in zebrafish mutants combining Csf1 ligands and receptors knockout will be instrumental in testing this hypothesis.

A major finding of our study is the demonstration that the second microglial wave in zebrafish is completely abolished in absence of *csf1rb,* thus uncovering a selective role for *csf1rb* in the establishment of HSC-derived adult microglia. Indeed, although microglia cells are present in each single mutant (albeit at comparably reduced cell density), lineage tracing of definitive microglia development revealed the microglial population presents in the brain of *csf1rb^-/-^* adult fish remain of primitive origin. This is in sharp contrast to *csf1ra*-deficient and *wild-type* fish, where the HSC-derived adult population fully replaces the primitive microglial pool. Interestingly, however, although adult microglia develop normally in *csf1ra*-deficient juvenile fish, their numbers drop towards the adult stage. This suggests that while being dispensable for the ontogeny of the definitive microglial wave, *csf1ra* likely contributes to microglia maintenance within the adult brain parenchyma. The opposite microglial phenotypes observed in *csf1ra* and *csf1rb* mutants also shed light on potential dynamics between the two microglial waves during development. Through timecourse analyses, we found that the incomplete recovery of primitive microglia in *csf1ra* juveniles is compensated by an earlier establishment of the definitive wave compared to *wild-type.* Conversely, the lack of definitive microglia in *csf1rb* mutants results in the primitive pool being retained in the adult. By analogy with the current view established in the mouse model (Guilliams et al., 2020), it is conceivable that competition for the juvenile brain niche regulates the exchange between the two microglial waves, in a scenario where efficient seeding of the brain by definitive microglia would require the regression of the primitive wave. Another plausible hypothesis is that the adult wave may actively participate to the removal of the primitive population, therefore explaining the maintenance of embryonic microglia in *csf1rb*-deficient adult animals. Future studies, making use of new and more sophisticated tools, will be required to address these complex questions. Nevertheless, since neither definitive microglia in *csf1ra* mutants nor primitive microglia in *csf1rb* mutants achieved the cellular density seen in *wild-type* adults, other intrinsic or extrinsic factors aside from niche availability are likely to affect microglia homeostasis in the adult brain.

Our investigations into the molecular mechanisms underlying the microglial phenotype of *csf1rb* mutant fish revealed that HSCs give rise to myeloid cells in a *csf1rb*-dependent fashion. Indeed, we found that *csf1rb*-deficient fish display a broad deficit in definitive myelopoiesis, as supported by the decrease of HSC-derived macrophages during embryonic development and in the adult hematopoietic niche. These findings are consistent with the *runx1*-dependent selective expression of *csf1rb* on blood progenitor cells throughout life. In addition, *csf1rb* is dispensable for HSC emergence in the AGM and, unlike *csf1ra,* does not seem to control cell migration to the different niches. On the whole, this suggests that the depletion of adult microglia and definitive macrophages in the *csf1rb* mutant results from a deficit of differentiation at the level of HSC-derived myeloid progenitors. These data add to previous findings in zebrafish (Yu et al., 2017) and mouse (Azzoni et al., 2018) showing that distinct molecular mechanisms regulate the emergence of subsequent macrophages waves. Overall, zebrafish *csf1ra* and *csf1rb* mutants may thus provide insightful models for the functional dissection of each microglial population and to better understand microglia development from an evolutionary perspective.

Finally, a surprising finding of our study is that fish lacking the *csf1rb* paralog also lack a population of *mpeg1^+^* B cells in the WKM. As we previously showed that these cells account for the majority of IgM-expressing B lymphocytes in zebrafish (Ferrero et al., 2020), these observations suggest that B lymphopoiesis is globally impaired in *csf1rb* mutant animals. This is interesting because while no such phenotype has been reported in adult CSF1R-deficient mice so far, expression of CSF1R was recently identified on a subset of embryonic myeloid-primed B-cell progenitors in the fetal liver, and its loss associated to defective fetal B-cell differentiation *in vivo* (Zriwil et al., 2016). Although the precise contribution of *csf1rb* to zebrafish B lymphopoiesis remains to be investigated, our work thus provides further support for a role of CSF1R beyond myelopoiesis in vertebrates. Also, because in the mouse the fetal wave of B lymphopoiesis mainly produces innate-like B-1 lymphocytes, and given the B-cell phenotype similarities between *Csf1r*-deficient mice and *csf1rb^-/-^* zebrafish, this study also adds to the growing view that mammalian B-1 lymphocytes and teleost adult B cells could be evolutionary related (Scapigliati et al., 2018).

### MATERIALS AND METHODS

#### Zebrafish husbandry

Zebrafish were maintained under standard conditions, according to FELASA guidelines (Alestrom et al., 2019). All experimental procedures were approved by the ethical committee for animal welfare (CEBEA) from the ULB. The following lines were used: *Tg(mpeg1:EGFP)^g/22^ (Ellett et al., 2011); Tg(kdrl:Cre)^s89^* (Bertrand et al., 2010); *Tg(actb2:loxP-STOP-loxP-DsRed^express^)^sd5^* (Bertrand et al., 2010); *panther^j4e1^* (Parichy DM, 2000); *csf1rb^sa1503^* mutants were generated via ethyl-nitrosurea (ENU) mutagenesis by the Sanger Institute Zebrafish Mutation Project. The following primers were used to identify the point mutation in intron 11 by PCR on genomic DNA: sa1503F *(5′-CTCTCTCTGTGGCAACTCTATGGATG-3′); sa1503R(5′-CGCTCCTGCTCCAAGAACCTG-3)*.

#### Flow cytometry and cell sorting

Single-cell suspensions of zebrafish whole embryos or adult WKM were prepared as previously described (Ferrero et al., 2018). Heads of 6 dpf zebrafish larvae were rapidly dissected in ice-cold PBS and then processed as the other samples. Flow cytometry and cell sorting were performed with a FACS ARIA II (Becton Dickinson). Analyses were performed using the FlowJo software (Treestar).

#### Quantitative PCR

RNA extraction from sorted cells and cDNA synthesis were performed as described (Ferrero et al., 2020). Biological triplicates were compared for each subset. Relative amount of each transcript was quantified via the ΔCt method, using *elongation-Factor-1-alpha (ef1α)* expression for normalization. Primers used are reported in Table 1.

**Table 1.**
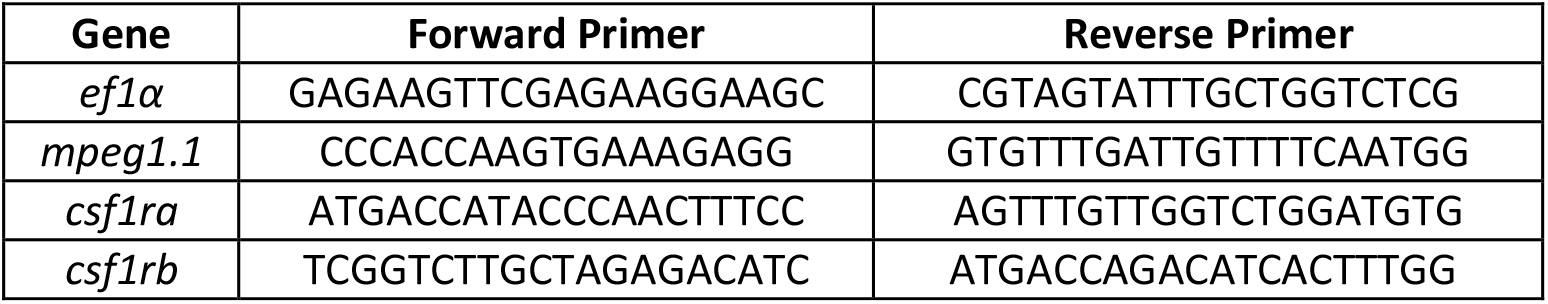
qPCR primers used throughout the paper

#### Whole Mount In Situ Hybridization (WISH)

Probes for *apoeb, runx, c-myb, mfap4* and *csf1rb* were synthesized *in-vitro.* For the *csf1rb* WISH, we combined two probes hybridizing to different portion of the transcript to increase the signal strength. The following primers were used for the generation of the *csf1rb* probes from zebrafish 4 dpf larvae cDNA: Fw1: 5′- ATCATTGCAGTGCTGACCTGTATG;Rv1:5′-GGTGAGCTCCAGGTGAAGTTGTAG; Fw2: 5^-/-^ATGGCCAACCAATCCATTTCTGAG Rv2: 5′-AGTAAGCATTCCTTGCGGGATGTT. Embryos or larvae were fixed in 4% PFA at 4°C O/N and then stored in methanol at −20°C. Whole-mount *in-situ* hybridization was performed according to previously published protocols (Thisse and Thisse, 2008).

#### Immunofluorescence, tissue clearing and imaging

Larvae were fixed in 4% PFA O/N at 4°C and stored in methanol at −20°C. Adult zebrafish brains were fixed 4 hours in 4% PFA at 4°C, incubated in 30% sucrose/PBS overnight and embedded in OCT (Leica) for cryosectioning. Immunofluorescence on whole embryos or 30μm-thick brain slices was performed as described (Ferrero et al., 2018), using chicken anti-GFP (1:500, Abcam), polyclonal rabbit anti-DsRed (1:500, Takara) primary antibodies and goat-anti chicken Alexa 488 (1:500, Abcam), donkey anti-rabbit alexa 594 (1:500, Abcam) secondary antibodies. Dissected brains from larval or juvenile fish (21 to 50 dpf) were fixed in 4% PFA at pH 8.5 to preserve endogenous fluorescence, and subsequently tissue-cleared using the CUBIC protocol (Susaki et al., 2015), as described (Ferrero et al., 2018). Imaging was performed on a Zeiss LSM 780 inverted microscope, using a Plan Apochromat 20x objective for adult sections and a LD LCI Plan Apochromat 25x water-immersion objective for wholemount embryos and tissue-cleared brains. Images of entire adult brain sections were obtained by combining 15 tiles, for a total area of 1.80 mm^2^.

## ACKNOWLEDGEMENTS

We thank Mireia Rovira, member of the Wittamer lab, for critical discussion and comments on the manuscript. We are also grateful to Marianne Caron for technical assistance and to Christine Dubois for help with flow cytometry.

## DATA AVAILABILTY

All datasets generated for this study are included in the manuscript/ Supplementary Files.

## AUTHORSHIP CONTRIBUTION

V.W. designed the research and directed the study. G.F., M.M and E.D. performed experiments. G.F. and V.W. wrote the manuscript with comments from all authors.

---

Figure S1

**Figure S1.**
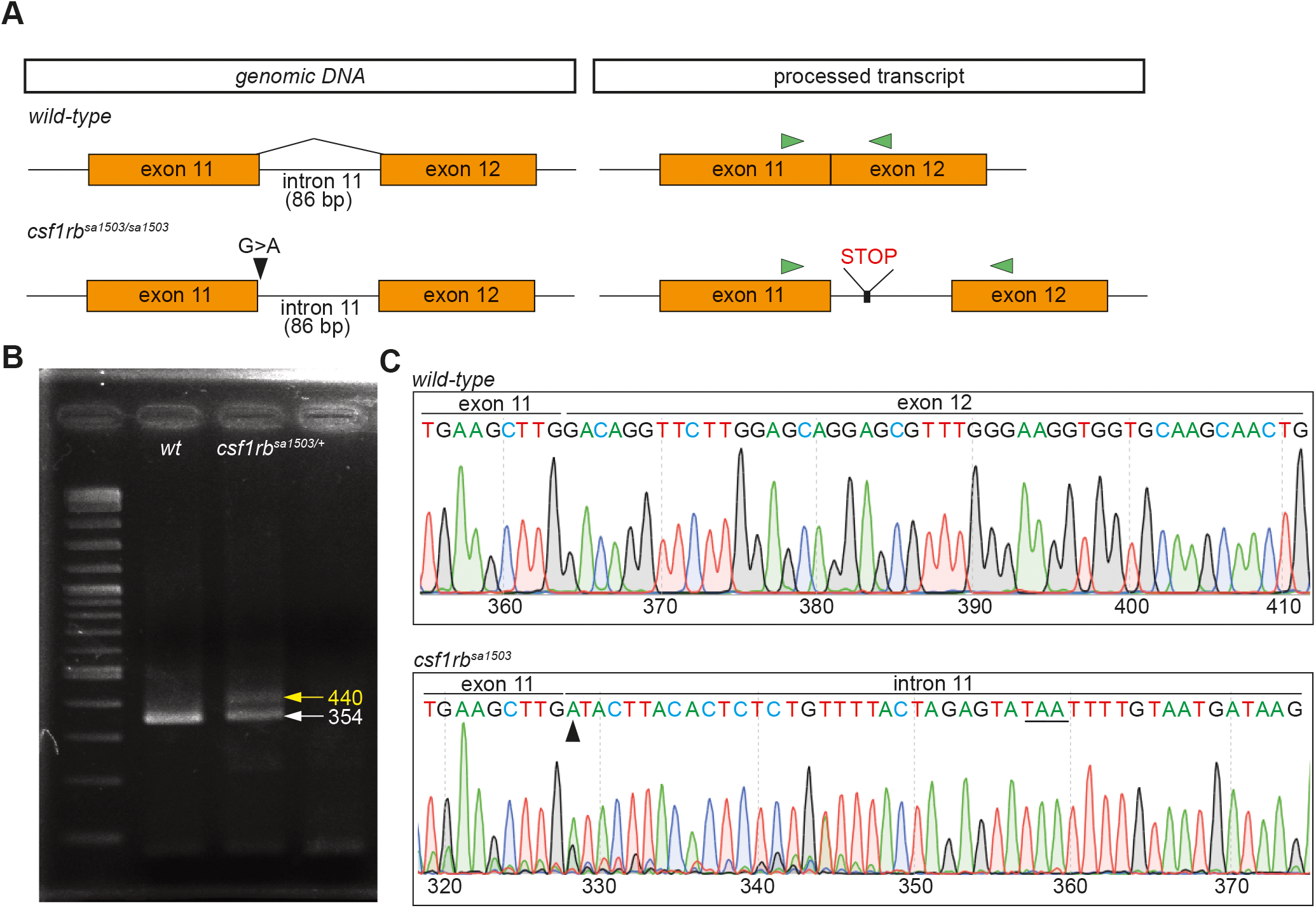
Characterization of the csf1rbsal5O3 mutant line. (A) Schematic view of the exons 11 and 12 of the *csf1rb* gene and the alteration caused by the splice mutation (black arrowhead) in the *csf1rb^sa1503^* line, which leads to the inclusion of 86-bp from intron 11 and the introduction of a premature stop codon in the coding sequence. Green arrowheads indicate the position of the primers used for PCR amplification (B) RT-PCR on whole brain isolated from adult *wild-type* and *csf1rb*^sa1503/+^ mutant, showing the spliced wild-type transcript at 354 bp and the unspliced *csf1rb*^sa1503^ transcript at 440 bp (+ 86 bp). As expected, both forms of the transcript are found in the heterozygous mutant. (C) Sequence chromatograms show the G>A substitution (black arrowhead) and the premature stop codon TAA (underlined) in the *csf1rb*^sa1503^ mutant.

Figure S2

**Figure S2.**
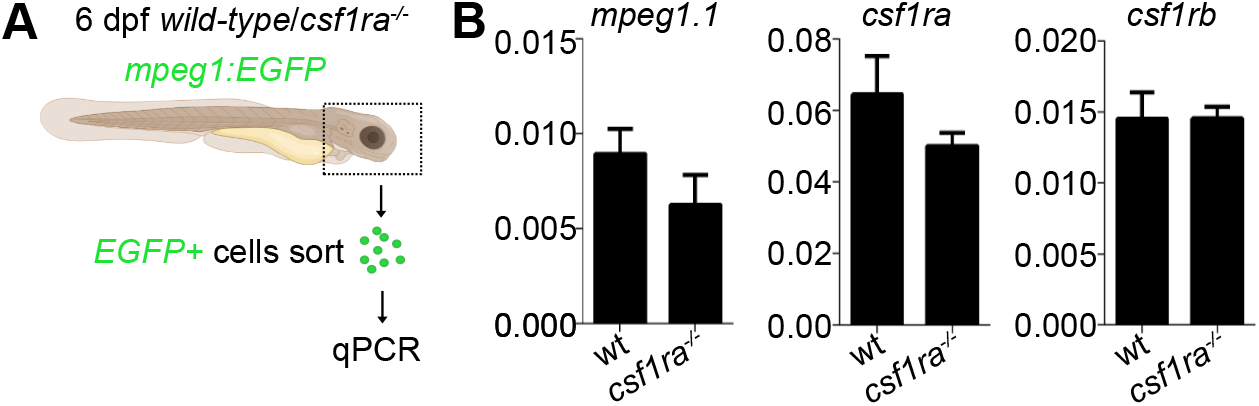
Comparison of csf1rb expression in 6 dpf *wild-type* and *csf1ra^-/-^* head macrophages. (A) Experimental outline. (B) qPCR analysis of gene expression for *mpeg1.1, csf1ra* and *csf1rb* in sorted *mpegl:EGFP^+^* cells. Error bars represent SEM (n=3). Values on the y-axis indicate transcript expression normalized to *eflα* expression level.

